# The Administration Effect of Lactic Acid Bacteria Reducing Environmental Alkyl and Fatty Acid Hydroperoxides on Piglets in the Absence of Antimicrobial Agents and in the Presence of Iron

**DOI:** 10.1101/2022.08.12.503799

**Authors:** Tatsuo Noguchi, Akio Watanabe, Yoshimasa Sagane, Kouji Nomoto, Junji Terao, Tomonori Suzuki, Masataka Uchino, Akira Abe, Youichi Niimura, Shuhei Ikeda

## Abstract

A lactic acid bacterium, *Lactiplantibacillus plantarum* P1-2 (*Lp*P1-2), can reduce environmental fatty acid hydroperoxides. The administration of *Lp*P1-2 to oxygen-sensitive short-lived nematode mutants and iron-overloaded rats reduced the oxidative stress-related index. Since young piglets have a weak defense system against oxidative stress and are vulnerable to environmental stress, antimicrobial agents have been administered in the rearing. Based on these results, we investigated the effect of *Lp*P1-2 administration to prepartum sows and infant piglets until weaning without antimicrobial agents on the growth of young piglets.

The group including both sows and piglets that were administrated with lactic acid bacteria containing *Lp*P1-2 (LAB*Lp*) until the end of lactation showed the growth-promoting effect of piglet from lactation to early regular rearing, and even in late regular rearing. Blood biochemical markers were in the normal ranges in both LAB*Lp*-administrated and non-administrated groups, but various disease-related markers tended to decrease in the administrated group.

To investigate the effects of LAB*Lp* administration on postpartum piglets, the piglets born from prenatally LAB*Lp*-administrated sows were divided into two groups and then administrated with or without LAB*Lp*. The piglets in the LAB*Lp*-administrated group tended to grow very slightly higher than those in the non-administrated group from lactation to early regular rearing. After that, the growth in both groups was almost the same. These results suggest that LAB*Lp* administration to prepartum sows is essential for the growth-promoting effect.

The postnatal LAB*Lp*-administrated piglets showed a lower serum lipid peroxidation index than the non-administrated piglets, and had higher numbers of lactic acid bacteria and bifidobacteria in feces at the end of LAB*Lp* treatment. In fattening performances, the LAB*Lp*-administrated group showed a significant improvement in meat quality.

We also discuss the growth and physiological effects by *Lp*P1-2 administration with iron on piglets because iron administration is another important issue in piglet farming.

## Introduction

Antimicrobial resistance (AMR) infections are widespread worldwide [1, 2], and to prevent AMR, it is essential to limit the use of minimum antimicrobial agents to the minimum necessary and to prevent the release of them into the environment. There are two types of antimicrobial agents for livestock: those that are mainly used for the treatment of diseases [3] and those that are mainly used for the growth promotion of livestock [4]. The addition of both antimicrobial agents to livestock feed has been in common, making livestock indigenous bacteria drug resistant. As a result, it is concerned that their release into the environment may lead to horizontal transmission to humans. To prevent this, risk management measures for antimicrobial feed additives have been carried out [5].

In pig growth, young piglets are sensitive to environmental stress caused by heat and temperature changes [6-8], and antimicrobial agents have been administrated to young pigs to prevent disease and promote growth [3, 4]. In young piglets, it is known that oxidative stress index increases due to environmental stress [8]. Since oxidative stress is a cause of various diseases, enhancement of the defense system against oxidative stress in young piglets is expected to prevent the diseases and promote growth during the juvenile stage.

The authors isolated a lactic acid bacterium, (*Lp*P1-2), from fermented food. This strain reduces environmental alkyl hydroperoxides and fatty acid hydroperoxides and converts both peroxides to the less toxic hydroxy form via two electron reduction [9]. Oral administration of *Lp*P1-2 to the oxygen-sensitive short-lived nematode mutant resulted in a significant expansion of its lifespan, suggesting that *Lp*P1-2 inhibits internal oxidative stress [9]. To specify the organs involved in this response, we performed a similar experiment on iron-overloaded rats in which lipid peroxidation was induced. The administration of *Lp*P1-2 showed a significant reduction in the peroxidation index value in the colonic mucosa of these rats [9]. In this report, we examined about the effect of *Lp*P1-2 administration to prepartum sows and postpartum piglets for the purpose of enhancing the defense power in young piglets against oxidative stress.

In piglet rearing, iron supplements are administered to piglets to prevent iron deficiency anemia during the growth process[10-12]. However, since iron is a factor that causes the Fenton reaction [13], piglets with immature oxidative stress defense systems may exhaust energy to cope with such stress. In this report, to develop a probiotic material to replace antimicrobial agents, we created an antimicrobial-free feed supplemented with iron and *Lp*P1-2, and investigated the effect of *Lp*P1-2 on the growth of young piglets.

## Materials and Methods

### Test animals

In Test I, two sows of WL breed (Yorkshire - Landrace) born in January 2009 were used as test sows and were synchronized to have the same farrowing date. One sow (α) was given lyophilized LAB*Lp* powder from 28 days before farrowing until weaning, and the other one (β) was given skimmed milk powder as a placebo (Fig 1). The sows were artificially inseminated with Duroc semen and the piglets born (27 in total) were used as test piglets. A mother-child exchange was made for half of the newborn piglets. The piglets were weaned 28 days after farrowing. Lyophilized LAB*Lp* powder was administered to the piglets suckled by the sow α (14 piglets, LAB group, piglets A (n=7) and piglets C, (n=7)) from immediately after delivery until weaning. Skimmed milk powder was administered to the piglets suckled by the sow β (13 piglets, Control group, piglets B (n=6) and piglets D, (n=7)) instead of lyophilized LAB*Lp* powder. No antimicrobial agents were added to the diets of either sow α or β. We monitored the weights gain and blood biochemistry of the piglets during rearing. In Test II, as in Test I, one WL (Yorkshire - Landrace) sow was artificially inseminated with Duroc semen, and the piglets (10 piglets) born were used as test piglets (Fig 2). The piglets were divided into two groups, half of which received lyophilized LAB*Lp* powder immediately after delivery until weaning (LAB group), and the other half were administered skimmed milk powder instead of lyophilized LAB*Lp* powder (Control group). The administration of LAB*Lp* to the sow and the weaning were the same as the sow α in Test I. The test period was from piglet farrowing to the end of fattening in both Tests I and II.

**Fig. 1.**
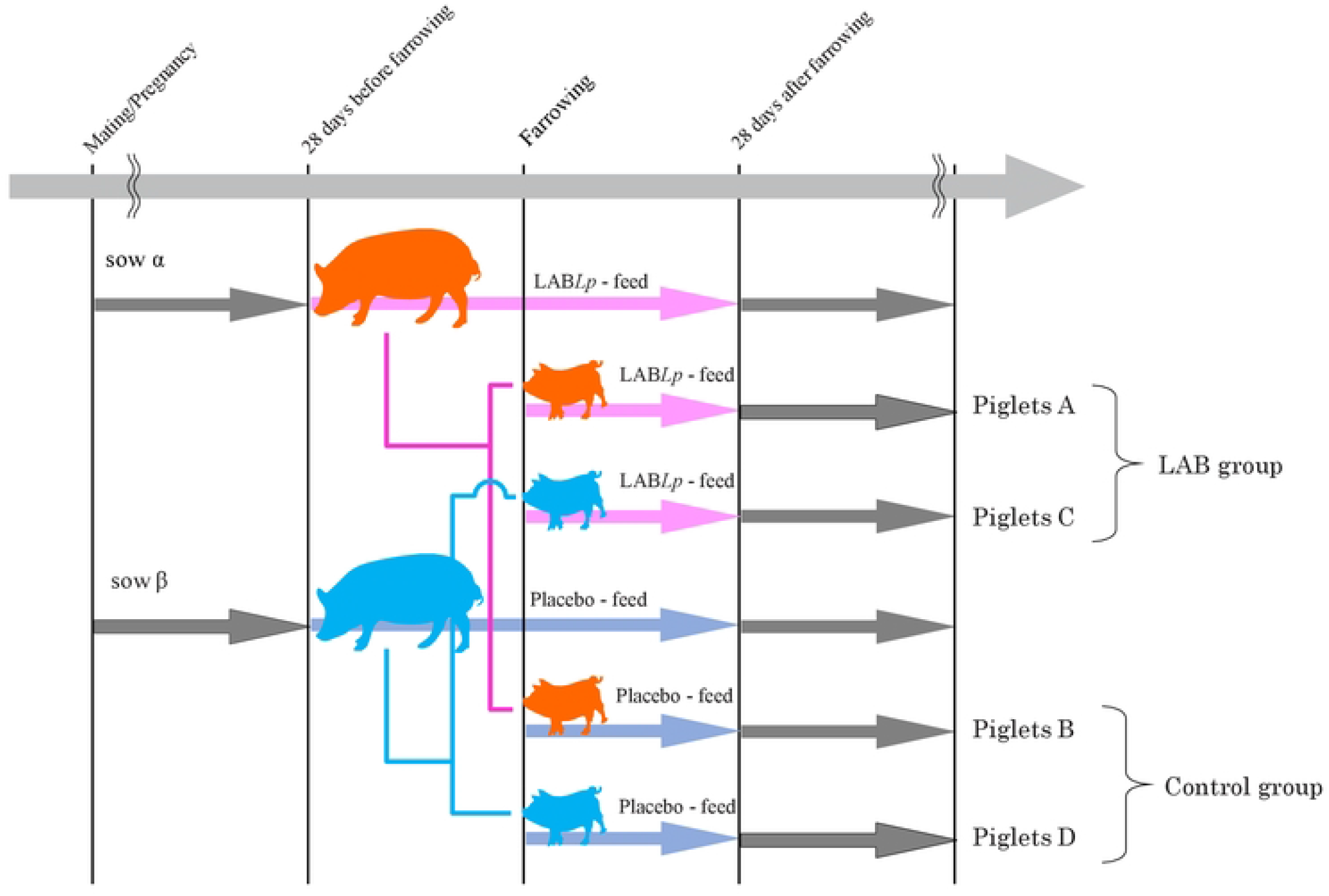
Schedule of Test I. The pink arrow represents lyophilized LAB*Lp* powder administration (LAB group, piglet A (n=7) and piglet C, (n=7)), and the blue arrow represents skimmed milk powder administration as a placebo (Control group, piglet B (n=6) and piglet D (n=7)). The light gray arrow indicates the administration of conventional feed without any antimicrobial agents, as described in the Materials and Methods. The mother sow and her biological piglets are shown using the same color.

**Fig. 2.**
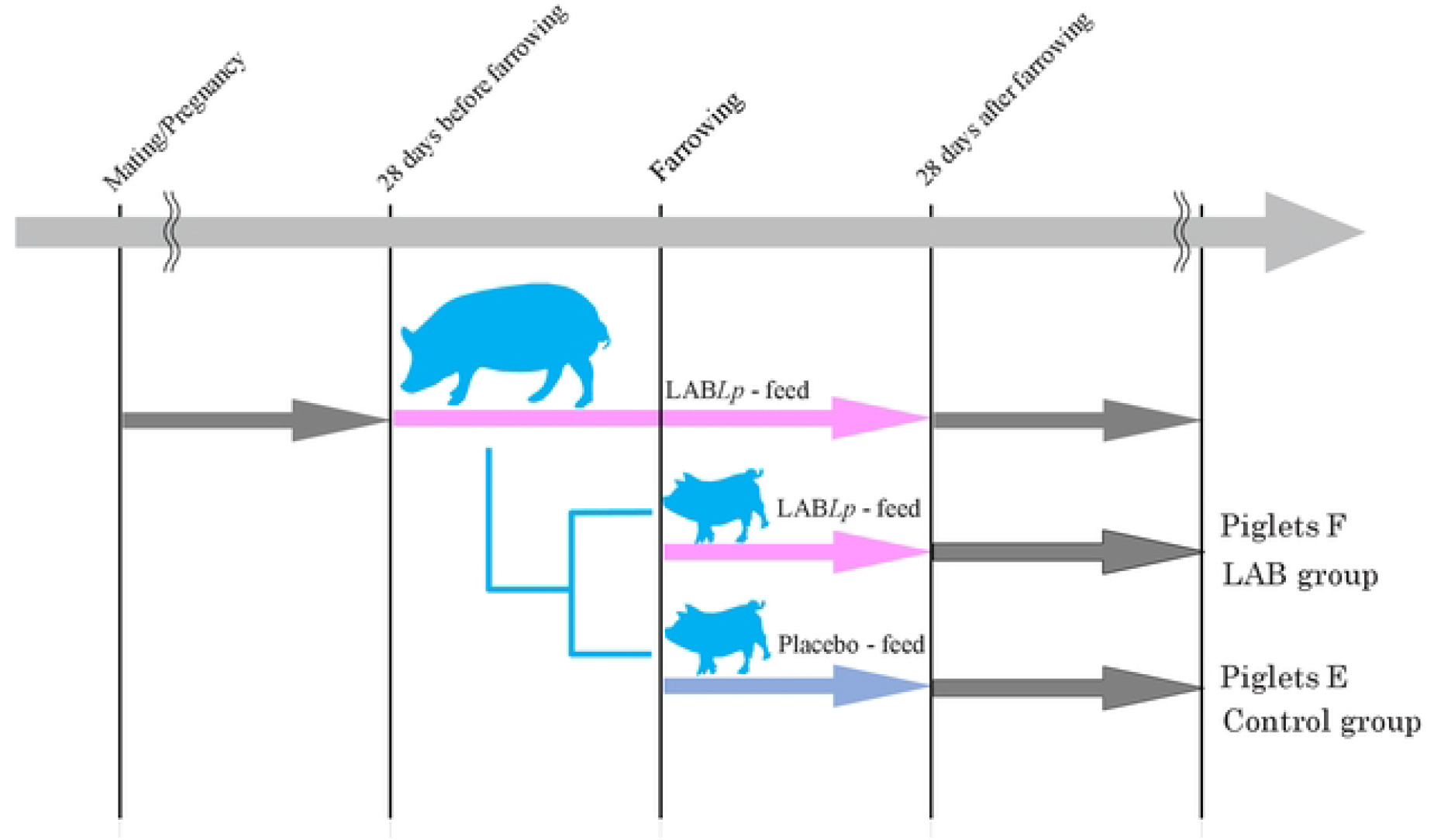
Schedule of Test II. The pink arrow represents lyophilized LAB*Lp* powder administration (LAB group, piglets F, n=5), and the blue arrow represents skimmed milk powder administration as a placebo (Control group, piglets E, n=5). The light gray arrow indicates the administration of conventional feed without any antimicrobial agents, as described in the Materials and Methods. The mother sow and her biological piglets are shown using the same color.

In Test II, the serum lipid peroxidation index, body weight gain, and fecal microbial flora of piglets were investigated. Piglets of finishing phase were examined for body weight, branch weight at slaughter, yield, back fat thickness, and meat quality when they were shipped at 157 days of age.

### Test feed

Each sow was fed 2.5 kg/day of sow formula (Big Mama, Toyohashi Feed Co., Ltd., Japan) containing no antimicrobial agents before farrowing and 5.0 kg/day after farrowing. The sow α was administrated 25 g of lyophilized LAB*Lp* before farrowing, and 50 g after farrowing before feeding. The sow β received 25 g of skimmed milk powder as a placebo before farrowing and 50 g after farrowing before feeding, instead of lyophilized LAB*Lp* powder. A homemade diet (Table S1) given to the piglets were gradually increased from birth to 56 days of age. After birth, 1 g of lyophilized LAB*Lp* dispersed in 10 ml of pure water per piglet was administrated to piglets A and C in the LAB group (Fig 1). In the control group of piglets B and D, 1 g of skimmed milk powder as a placebo was administered instead of the lyophilized LAB*Lp* powder (Fig 1). For iron supplementation, 800 mg/ml of iron dextran (equivalent to 200 mg/ml of iron, Iron Syrup S, Scientific Feed Laboratory Co., Ltd., Japan) was orally administrated at a dose of 1 ml/head on the third day after birth. Other rearing management followed the conventional rearing method of Fuji Farm, Tokyo University of Agriculture.

### Lyophilized lactic acid bacteria powder

In the preparation of lyophilized LAB*Lp* powder, LAB*Lp* was collected by centrifugation from a liquid medium consisting of 3% glucose, 3% yeast extract (Aromild, KOHJIN Life Sciences Co. Ltd., Japan), and 3% yeast extract (SK, Nippon Paper Group, Japan), mixed with 10% skimmed milk powder solution as a protective agent, and then freeze-dried. The ratio of the dry weight of LAB*Lp* to the weight of skimmed milk powder was adjusted to 1:9. The prepared LAB*Lp* powder was mixed with the basal diet and stored at -18°C until use. The *Lp*P1-2 in the LAB*Lp* powder after preparation was 1×10^10^ cfu/g.

### Measurement of the number of lactic acid bacteria and bifidobacteria in feces

BL medium, modified LBS medium, BS medium, and NN medium prepared by the method of Mitsuoka *et al*. were used [14, 15]. Feces obtained from sows and piglets were suspended in diluted solution for dispersion, smeared on each medium agar plates, and the colonies that appeared were measured. The number of colonies obtained from BL medium, modified LBS medium and BS medium was counted as the number of total bacteria, lactic acid bacteria and bifidobacteria, respectively. The number of lecithinase-positive colonies obtained from NN medium was counted as the number of *Clostridium perfringens*.

### Assessment of serum lipid peroxidation

Lipid peroxidation markers such as malondialdehyde (MDA) in serum obtained from piglets at 28 and 56 days of age were measured by fluorescence using thiobarbituric acid reactive substances (TBARS) assay system (excitation wavelength at 515 nm, emission wavelength at 535 nm) [16, 17]. Serum protein concentration was determined by the method of Bradford [18]. The lipid peroxidation index was calculated as the TBARS value per serum protein.

### Blood biochemistry and blood cell count

Blood was collected from 28-day and 56-day-old piglets in groups A and D. For mother sows □ and □, it was done before LAB*Lp* administration and 28 and 56 days after delivery. Total protein, total bilirubin, albumin, aspartate aminotransferase (AST), alanine aminotransferase (ALT), alkaline phosphatase (ALP), lactic acid dehydrogenase (LDH), amylase, lipase, total bile acid (TBA), blood urea nitrogen (BUN), creatinine, total cholesterol (t-CHO), triglycerides (TG), inorganic phosphorus(P), chloride (Cl), sodium (Na), potassium (K), calcium (Ca), and serum blood glucose were investigated. Biochemical analysis in blood was performed using an automated analyzer (Fuji DRI-CHEM Fujifilm VET Systems, Japan) according to the JSCC standard method. For blood cell counting, an automated animal hematology analyzer PCE-170 (ERMA Sales Co., Ltd., Japan) was used.

### Meat quality evaluation

Carcass weight, backfat thickness and meat grading were estimated by the Japan Meat Grading Association (Tokyo 101-0063, http://www.jmga.or.jp) in according to the Pork Carcass Trading Standards, which evaluates the meat quality of pork in Japan.

### Statistical Analysis

The differences between two groups were analyzed by unpaired *t-*test. Data obtained from three or more groups were analyzed using non-repeated analysis of variance (non-repeated ANOVA). When the result of non-repeated ANOVA was significant, the Student–Newman–Keuls method was conducted. *P*-values of 0.05 or less were considered to have a significant difference.

## Results

### Test I

In two littermate sows (α and β) synchronized delivery, the sow (α) was administrated with lyophilized LAB*Lp* powder from 4 weeks before farrowing until weaning (28 days after farrowing). On the other hand, the sow (β) was administrated with skimmed milk powder (Fig 1).

Thirteen piglets (piglets A and C) were born from the sow α and 14 piglets (piglets B and D) from the sow β (Table S2). To evaluate the effect of LAB*Lp* administration before and after parturition, half of piglets born from each sow were suckled by mother sow exchange. Mother sow α fed 7 piglets born from her (piglets A) and 7 piglets born from mother sow β (piglets C). Mother sow β fed 7 pigs born from her (piglets D) and 6 pigs born from mother sow α (piglets B). Piglets A and C were administrated with lyophilized LAB*Lp* powder from immediately after farrowing until weaning. Piglets B and D were administrated with milk powder as a placebo instead of lyophilized LAB*Lp* powder. Under these rearing conditions, the delivered piglets were weighed (Fig 1, Table S2).

From the lactation period (28 days after birth) to the early normal rearing period (28 to 56 days of age), the weight gain after 7 days of age in Group A (piglets A) was higher than that in Group D (piglets D), and even in the late normal rearing (56 to 180 days of age) (Fig 3). The weight gain in Group B (piglets B) was lower than that in Group D (S1 Fig). The weight gain in Group C (piglets C) was almost the same as that in Group D (S1 Fig). In Groups A and D, the mother-child system at birth was continued, while in Groups B and C, mother-child exchange was carried out after delivery. The difference in weight gain observed between Groups B and C might be due to the stress on the piglets caused by mother-child exchange.

**Fig. 3.**
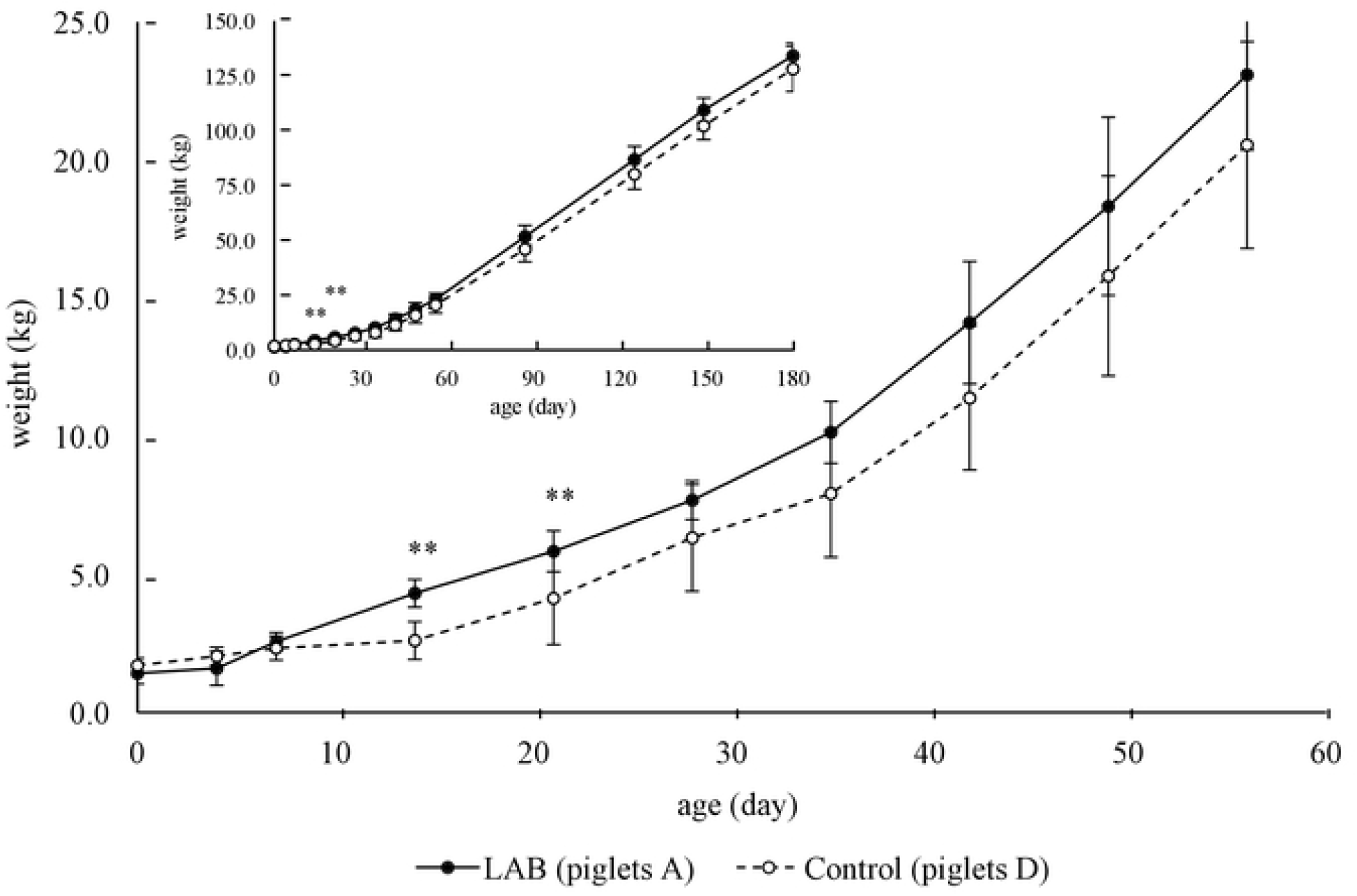
Growth curve of piglets in Test I. The solid line with black circle indicates the average weight of LAB group (piglets A (n=7) in Fig. 1), and the dotted line with white circle indicates the average weight of Control group (piglets D (n=7) in Fig. 1). The data is expressed as the mean value ± standard deviation (SD). Double asterisks (**) indicate a significant difference between the two groups (*p* < 0.01, unpaired *t*-test). The inset shows the weight change during the entire rearing period.

Subsequently, to see the effect of LAB*Lp* on the growth of piglets, a biochemical comparison of the blood of piglets A (the offspring of sow α, LAB*Lp* administration before and after weaning) and D (the offspring of sow β, non-administration before and after weaning) without mother-child exchange was carried out.

In both groups, all 24-test items remained within the normal range, regardless of the presence or absence of LAB*Lp* administration (Fig 4, S2 Fig, S3 Fig). The tests in which significant differences were observed between the two groups are described below.

**Fig. 4.**
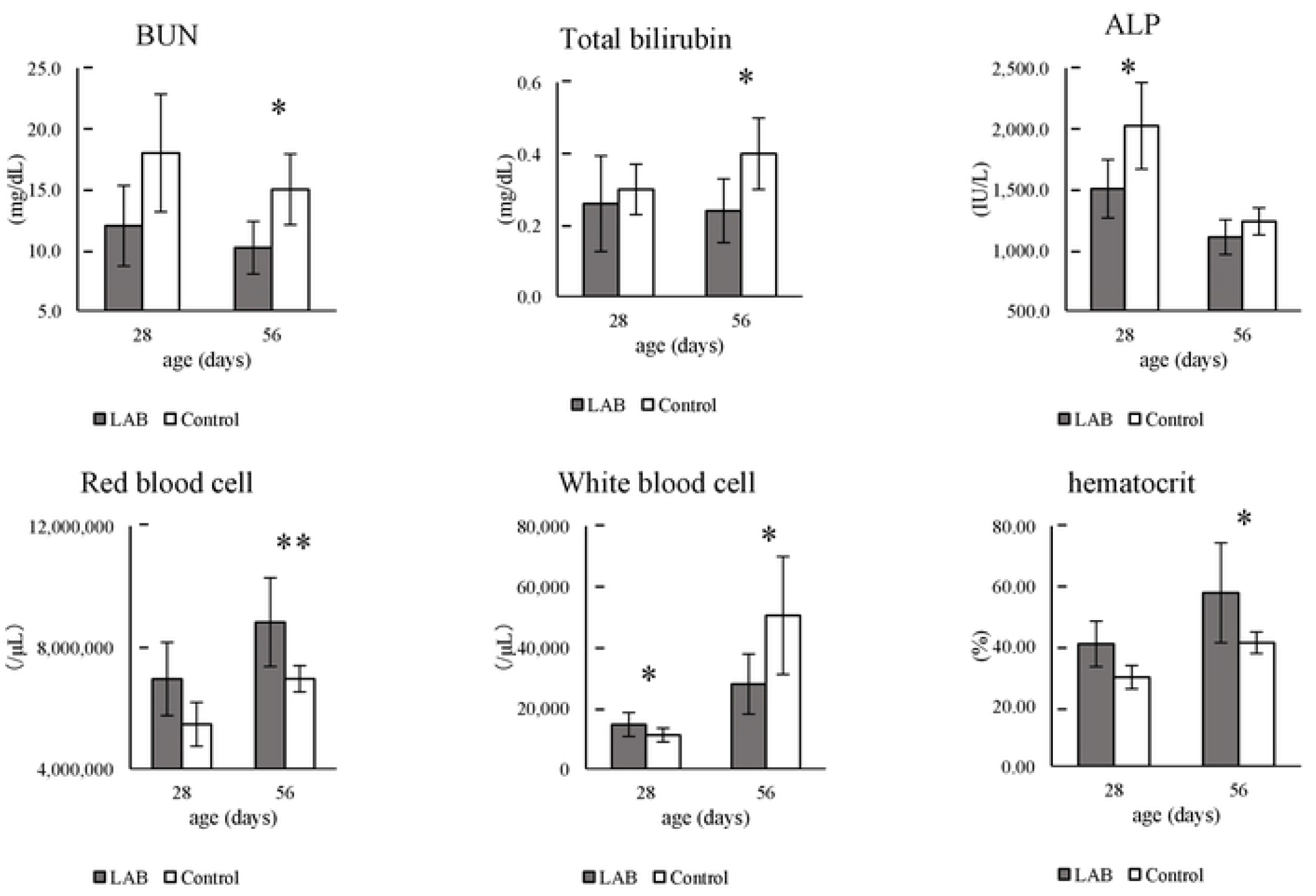
Blood biochemistry, blood cell counting of piglets in Test I. The gray bar indicates the average value of piglets A of LAB group (n=5), and the white bar indicates the average value of piglets D of Control group (n=5) in Fig. 1. The data is expressed as the mean value ± standard deviation (SD). An asterisk (*) and double asterisks (**) indicate a significant difference between the two groups (*p* < 0.05 and *p* < 0.01, respectively, unpaired *t*-test).

The number of red blood cells and hematocrit values were significantly higher in the LAB*Lp* - administrated (administration) group than in the non-administrated (control) group at 56 days of age. On the other hand, the white blood cell count, which is a marker that increases with bacterial infection, was significantly higher in the administration group than in the control group at 28 days of age. However, it reversed at 56 days of age, and the control group showed higher values.

ALP is a marker for liver, hepatic pathway, small intestine, and bone disorder, total bilirubin level is a marker for liver disorders, and BUN is a marker for renal disorders. ALP was higher at in the control group than in the administration group at 28 days of age. Total bilirubin and BUN were higher in the control group than the administration group at the age of 56 days.

In Group A, in which the mother sow was administrated with LAB*Lp* before and after delivery, a growth promoting effect on the piglets was observed not only during the lactation period but also during the early normal rearing period (28-56 days of age) after the end of administration (Fig 3). The detection values of the above disease markers in the blood during the normal rearing period were lower in the administration group than in the control group. The growth-promoting tendency in the administration group was also observed during the late normal rearing period (56-180 days of age) (Fig 3).

### Test II

The following experiment was conducted to examine the effect of LAB*Lp* administration after delivery. To avoid the stress caused by mother-child exchange, piglets born from the mother sow (sow □ in Fig 2) which was administrated with lyophilized LAB powder before delivery were divided into two groups, LAB*Lp* administrated group (Piglets F, in Fig 2) and non-administrated group (Piglets E, in Fig 2), and the effect of LAB*Lp* administration after delivery was investigated (Fig 2, Table S2).

The growth of piglets before and after weaning in the LAB*Lp* -administrated group F tended to increase slightly more than that in the non-administrated group E, but it was not significant, and then the growth in both groups was almost the same (Fig 5). The change in body weight in both groups was almost the same as that of the LAB*Lp*-administrated group (LAB in Fig 3) consisting of the piglets born and raised from the sow α in Test I.

**Fig. 5.**
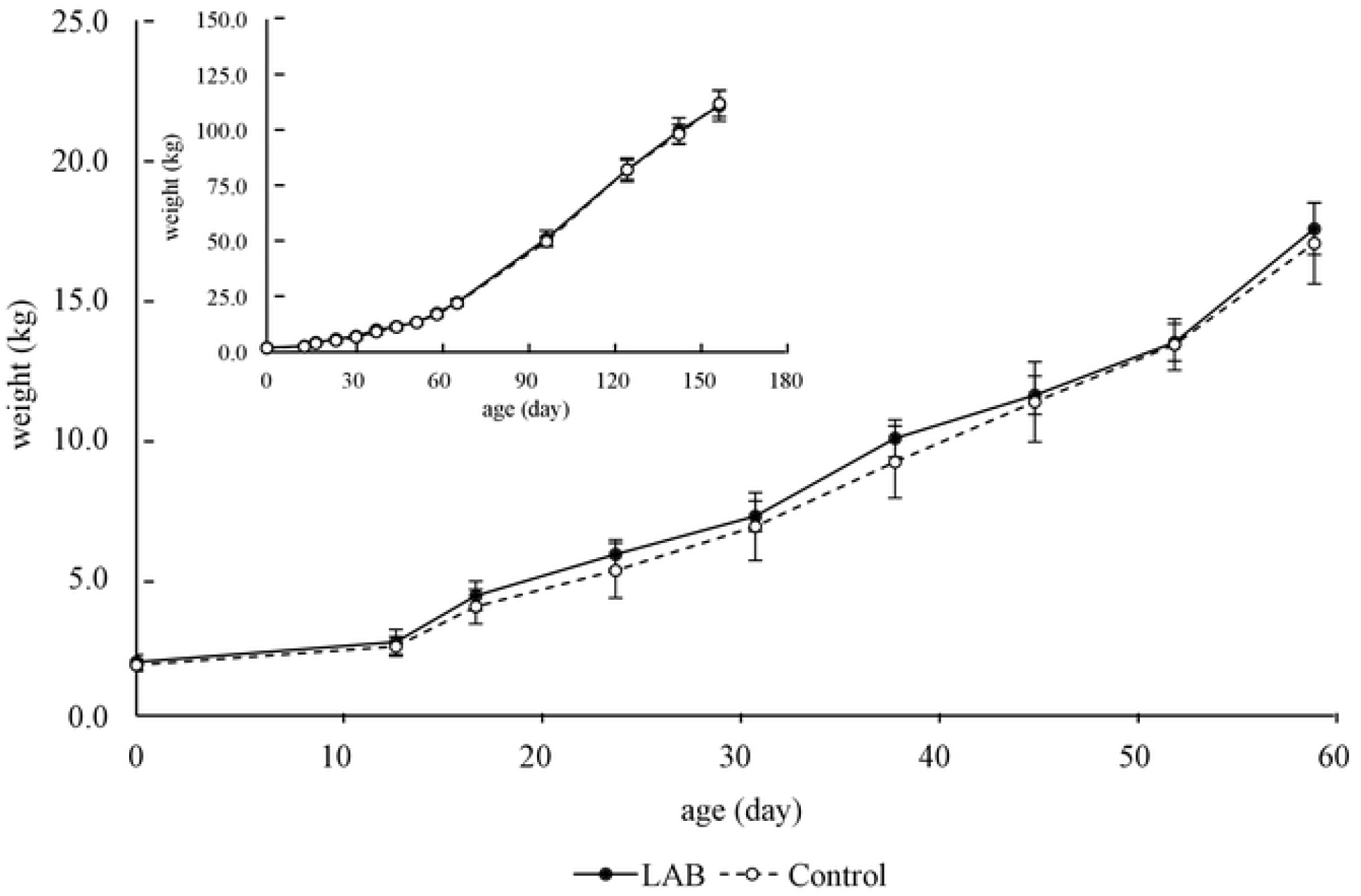
Growth curve of piglets in Test II. The solid line with black circle indicates the average weight of LAB group (piglets F, n=5) in Fig. 2, and the dotted line with white circle indicates average weight of Control group (piglets E, n=5) in Fig. 2. The data is expressed as the mean value ± standard deviation (SD). The inset shows the weight change during the entire rearing period.

The lipid peroxidation index in the serum of the LAB administrated group F was lower than that of the control group E at both 28 and 56 days of age (Fig 6). The numbers of both lactic acid bacteria and bifidobacteria in feces were high at weaning (28 days of age) after completion of LAB*Lp* administration. The number of lactic acid bacteria count was also high in the early weaning period (56 days of age) after completion of LAB*Lp*, but the number of bifidobacterium count was the same as that of the control group at 56 days of age. The number of *Clostridium perfringens* was not different between the two groups and was below the detection limit at 56 days of age (Fig 7). After finishing normal rearing (26 weeks of age), in piglet fattening performance the LAB-administrated group showed better results than those of the non-administrated group in terms of carcass weight, yield, grade, and significant improvement in meat quality (Table 1).

**Table 1.** Meat quality evaluation of shipped piglets in Test II. Results of meat quality evaluation of LAB group piglets F (n=5) and Control group piglets E (n=5) in Fig. 2 at the time of shipment. Carcass weight, backfat thickness and meat grading were estimated according to the Pork Carcass Trading Standards of the Japan Meat Grading Association (JMGA), which evaluates the meat quality of pork in Japan. The yield (%) in the table is defined as 100 x (weight at shipping) / (carcass weight). The data is expressed as the mean value, ± standard deviation (SD). An asterisk (*) indicates a significant difference between the two groups (p < 0.05, unpaired t-test).

| group | weight at shipping<br>(kg) | carcass weight<br>(kg) | yield<br>(%) | backfat chickness<br>(cm) | grade<br>(number of heads) |  |  |
| --- | --- | --- | --- | --- | --- | --- | --- |
|  |  |  |  |  | Excellent | Common | Out of grade |
| LAB | 111.0±6.8 | 74.1±4.1 | 66.76*±1.6 | 1.8±0.4 | 3 | 2 | 0 |
| Control | 112.0±6.0 | 71.5±3.6 | 63.84 ±1.8 | 1.9±0.6 | 1 | 3 | 1 |

**Fig. 6.**
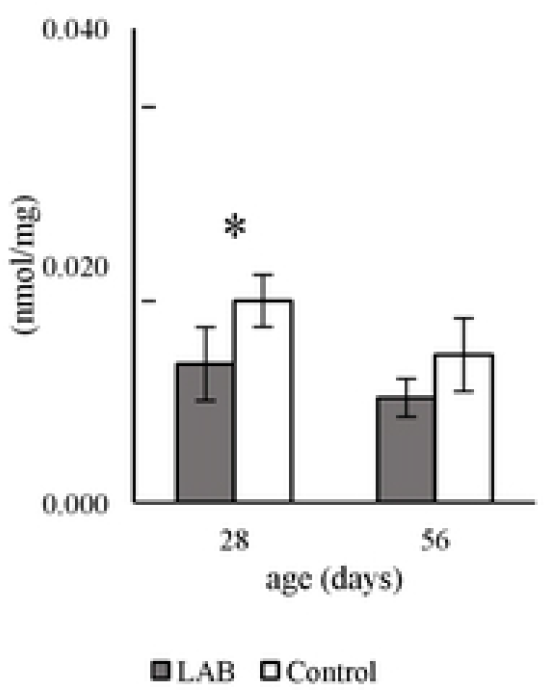
Lipid peroxidation in piglet serum in Test II. MDA, lipid peroxidation marker, was measured. The MDA levels are shown as TBAS values per serum protein level. The gray bar indicates the average value of piglets F of LAB group (n=5) in Fig. 2, and the white bar indicates average value of piglets E of Control group (n=5) in Fig. 2. The data is expressed as the mean value, ± standard deviation (SD). An asterisk (*) indicates a significant difference between the two groups (*p* < 0.05, unpaired *t*-test).

**Fig. 7.**
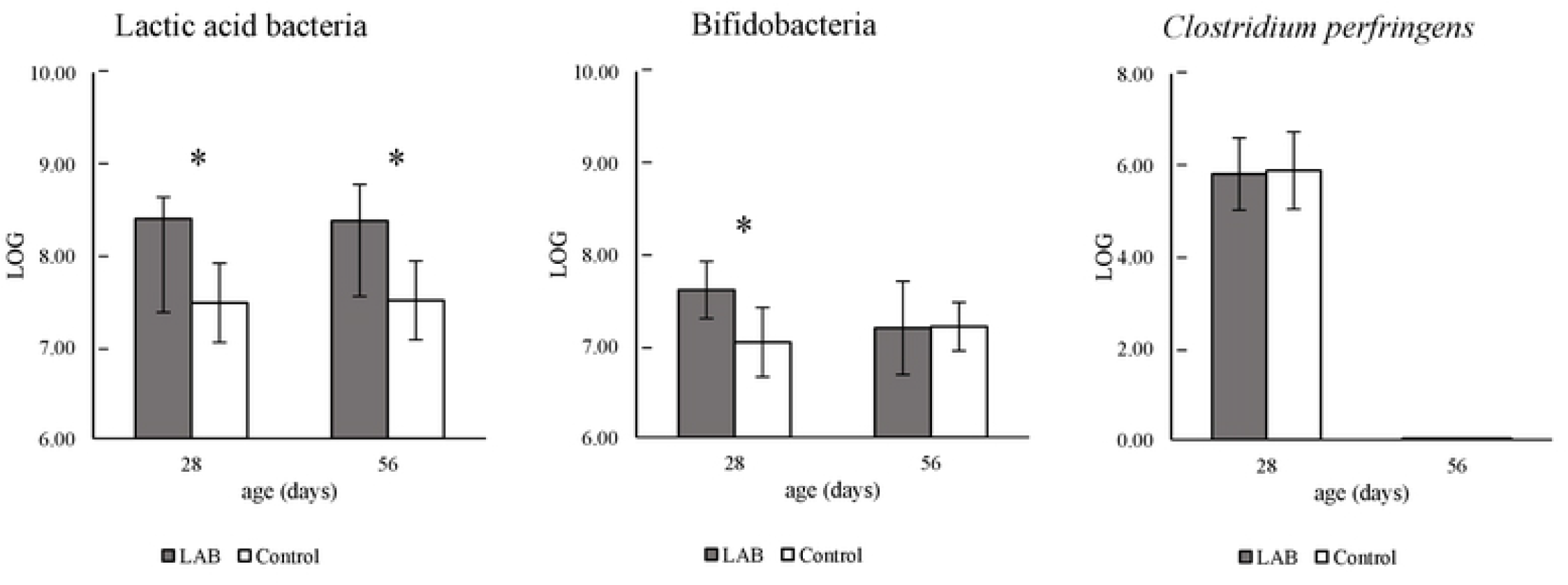
Microbial profile in piglet fecal samples in Test II. Feces obtained from piglets each group, LAB group and Control group in Fig. 2, were suspended in diluted solution for dispersion, smeared on agar plates as described in the Materials and Methods. The colonies that appeared on the plates were measured as described in the Materials and Methods. Results are presented as the mean (LOG: logarithm) ± standard deviation (SD). An asterisk (*) indicates a significant difference between the two groups (*p* < 0.05, unpaired *t*-test).

## Discussion

The LAB*Lp*-administrated group of mother sow and her delivered piglets showed slightly higher growth promotion (weight gain) in the young piglets than the non-administration group of those. In addition, the piglets born from the LAB*Lp*-administrated mother pig had a similar growth-promoting effect without LAB*Lp* administration after birth. This suggests that the administration of LAB*Lp* to sows before delivery is the key to growth promotion. On the other hand, the administration of LAB*Lp* to piglets after delivery showed a decrease in lipid peroxidation markers, an increase in bifidobacteria and lactic acid bacteria in feces, and significant improvement in meat quality. These physiological effects observed in the absence of antimicrobial agents indicate that lactic acid bacteria having reducing activity of fatty acid hydroperoxide shows an alternative effect of antimicrobial agents. The alternative effect of antimicrobial agents is an important issue in the field of livestock probiotics, and vigorous research on lactic acid bacteria including the same genus and species as this strain has been conducted [19-31]. In this study, not only the alternative effect of antimicrobial agents but also additional physiological effects were observed. The following are our thoughts on these effects found in this study.

### I. Antibiotic Substitute Effect

Young pigs easily develop diseases due to environmental stress (heat and temperature changes) [6-8]., and the administration of antimicrobial agents has been tried to prevent such diseases [3, 5]. An increase in oxidative stress index has been observed at the time of exhaustion by environmental stress [8]. Oxidative stress is a cause of various diseases, and it is expected that strengthening the ability to suppress oxidative stress have a preventive effect on diseases in infancy.

In this study, we found a decrease in lipid peroxidation index by TBARS assay of blood collected from LAB-administrated piglets. Previously, we showed that the reduction of lipid peroxidation index and the suppression of oxidative stress by the administration of *Lp*P1-2 in iron-overloaded rats and oxygen-sensitive short-lived nematode mutants [9], suggesting that the administration of *Lp*P1-2 to infant pigs after birth also suppresses oxidative stress. It has been reported that the oxidative stress defense system in infant pigs is weak [32]. Probably, the administration with *Lp*P1-2 to infant piglets could evoke an increase in anti-oxidative stress capacity and as a result prevent various diseases caused by oxidative stress. Therefore, we believe that lipid peroxide-reducing bacteria such as *Lp*P1-2 have efficiency as an alternative to antibiotics.

### II. Physiological effect

Young piglets are easy to develop iron deficiency anemia, and iron administration has been tried to prevent this [11]. However, iron is a factor that causes the Fenton reaction [13], and iron administration cause energy depletion in piglets with immature oxidative stress defense systems [32]. Therefore, setting the iron dosage according to the rearing condition of piglets is a very important issue in iron-administered rearing [10, 11, 33-36].

Previously, a decrease in lipid peroxidation index was observed in the colonic mucosa of iron-overloaded rats administrated with *Lp*P1-2 [9]. In the present study, TBARS level in serum samples iron-fed piglets in the LAB*Lp*-administrated group was lower than that in the non-administrated group. At the same time, the LAB*Lp*-administrated group showed a tendency to a decrease in the cytotoxicity index and growth-promoting effect, suggesting that the administration of *Lp*P1-2 suppresses iron toxicity over wide area in the piglet body. Possibly, as a result of the widespread suppression of iron toxicity in the iron-administrated piglets, the effect of iron as a supplement on iron deficiency anemia could be enhanced, and the nutritional status be also improved. Taken together, the improvement in nutritional status in the presence of iron supplement and the effect on disease prevention by the LAB*Lp* administration may be synergetic and contribute to the marked improvement in meat quality. Improvement of meat quality was also observed in the okara (soymilk by-product) fermented with immunobiotic lactobacilli [37]. In the future, it is necessary to analyze the mechanism of the meat quality improvement effect by probiotics.

In the research of probiotic lactic acid bacteria in pigs, there are some reports on the protective activity to oxidative stress [30, 38], but there is no report referring to the effect of lactic acid bacteria on iron toxicity. Although at this stage the mechanism by which *Lp*P1-2 suppresses iron toxicity is unknown, iron-dependent cell death (ferroptosis) has been reported to be involved in various human diseases [39-41]. In addition, the degradation of lipid peroxide [42] and the suppression of lipid peroxide toxicity [43-44] have been pointed out to be involved in the suppression of ferroptosis. Since *Lp*P1-2 has potent peroxidized fatty acid reducing activity, its administration may have suppressive effect on iron toxicity. Previously, we showed that *Lp*P1-2 must be administered as viable bacteria to exert this activity effectively [9]. The present study demonstrated its effectiveness in pigs. The number of *Lp*P1-2 found in the feces of the LAB-administrated group showed a higher than that of the non-administrated group (Fig 7). However, the present study did not directly prove the presence or absence of *L*pP1-2 in the feces during normal times. Analysis of the intestinal flora will be an issue for the future.

We expect that the inhibitory effect of lactic acid bacteria on the toxicity of administered iron observed in this study could be applied not only to reduce iron toxicity in pigs but also to inhibit ferroptosis. In the future, probiotics need to be verified from a wide perspective, containing analysis of the mechanism by which lactic acid bacteria such as *Lp*P1-2 suppress iron toxicity.

## Acknowledgements

We thank Mituo Sato of Tokyo University of Agriculture for helpful discussion, and Satoru Furukawa of the FRC for his technical advice. We greatly appreciate Prof. Soichi Arai at Tokyo University of Agriculture for valuable advice. This work was supported by grants from the Japanese Ministry of Education, Culture, Sports, Science and Technology.

## Supporting information

**S1 Fig.**
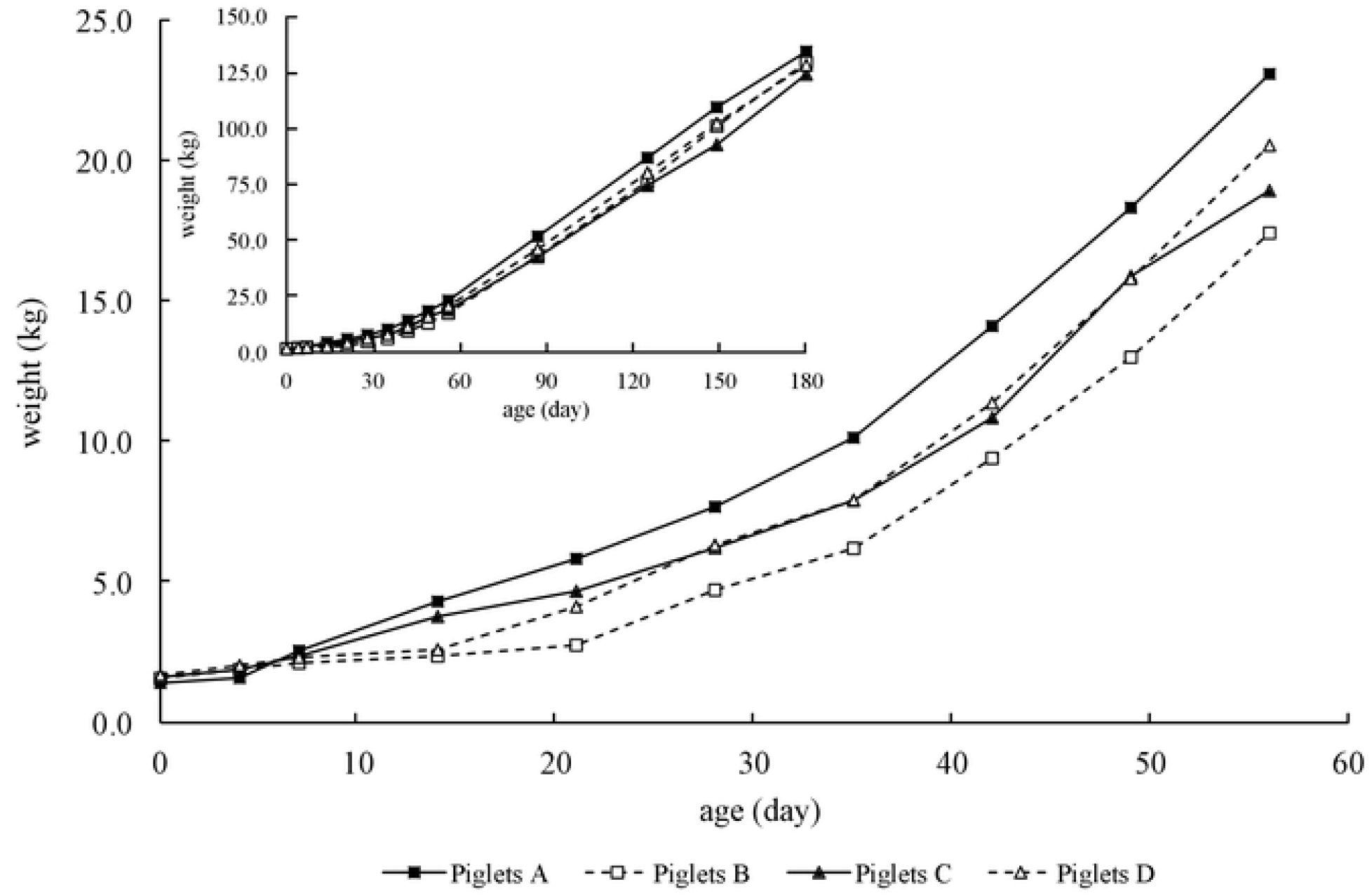
Growth curve of each litter piglet in Test I. The weight change in each litter piglet is shown in the figure. The solid line with black-square indicates the average weight of piglets A (LAB group (n=7) in Fig. 1) and the solid line with black triangle is the average weight of piglets C (LAB group (n=7) in Fig. 1). Also, the dotted line with white-square is the average weight of piglets B (Control group (n=6) in Fig. 1) and dotted line with white triangle is the average weight of piglets D (Control group (n=7) in Fig. 1). The inset shows the weight change during the entire rearing period.

**S2 Fig.**
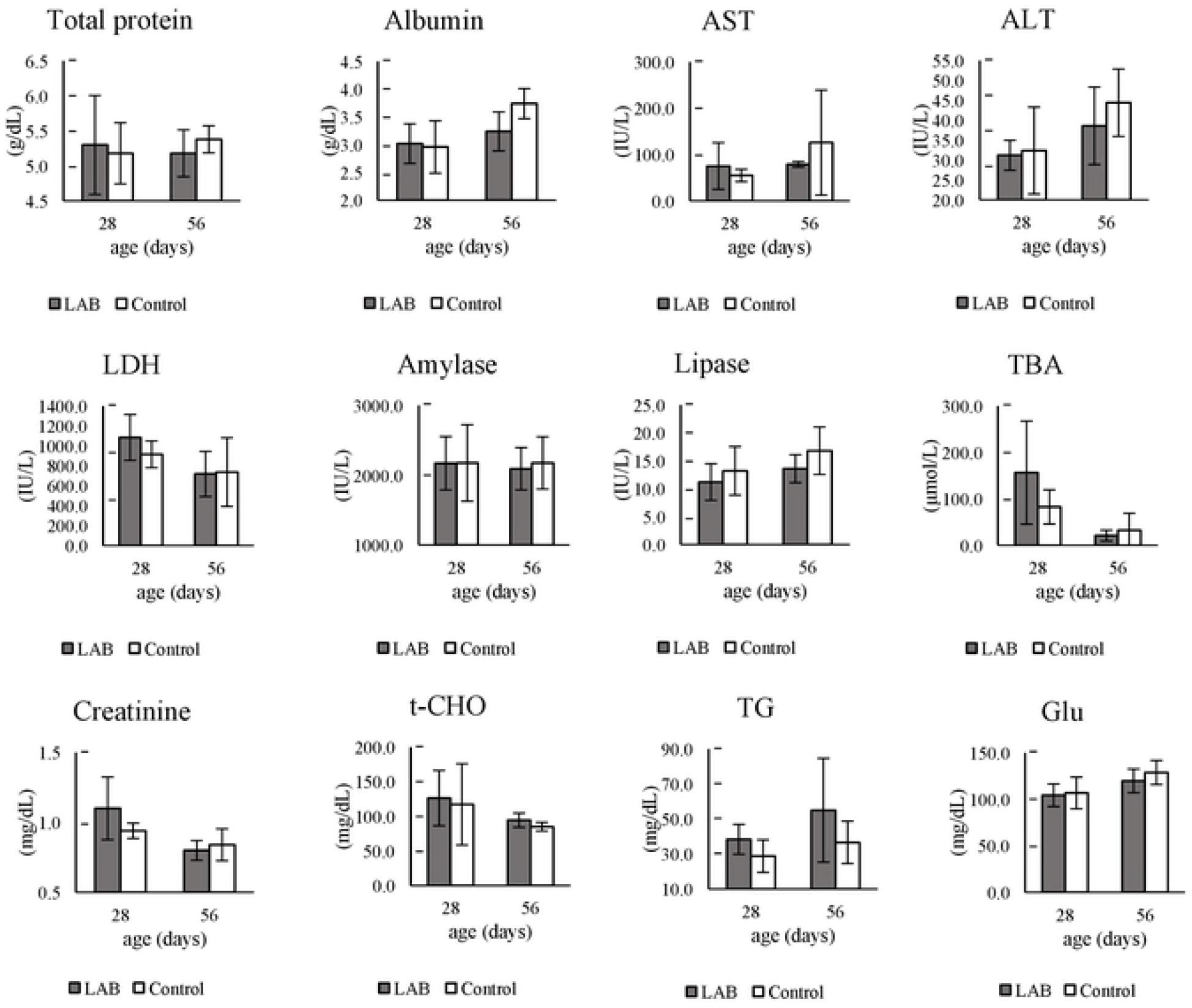
Blood biochemistry, blood cell counting of piglets in Test I. The gray bar indicates the average value of piglets A (LAB group (n=5) in Fig. 1), and the white bar indicates the average value of piglets D (Control group (n=5) in Fig. 1). The data is expressed as the mean ± standard deviation (SD).

**S3 Fig.**
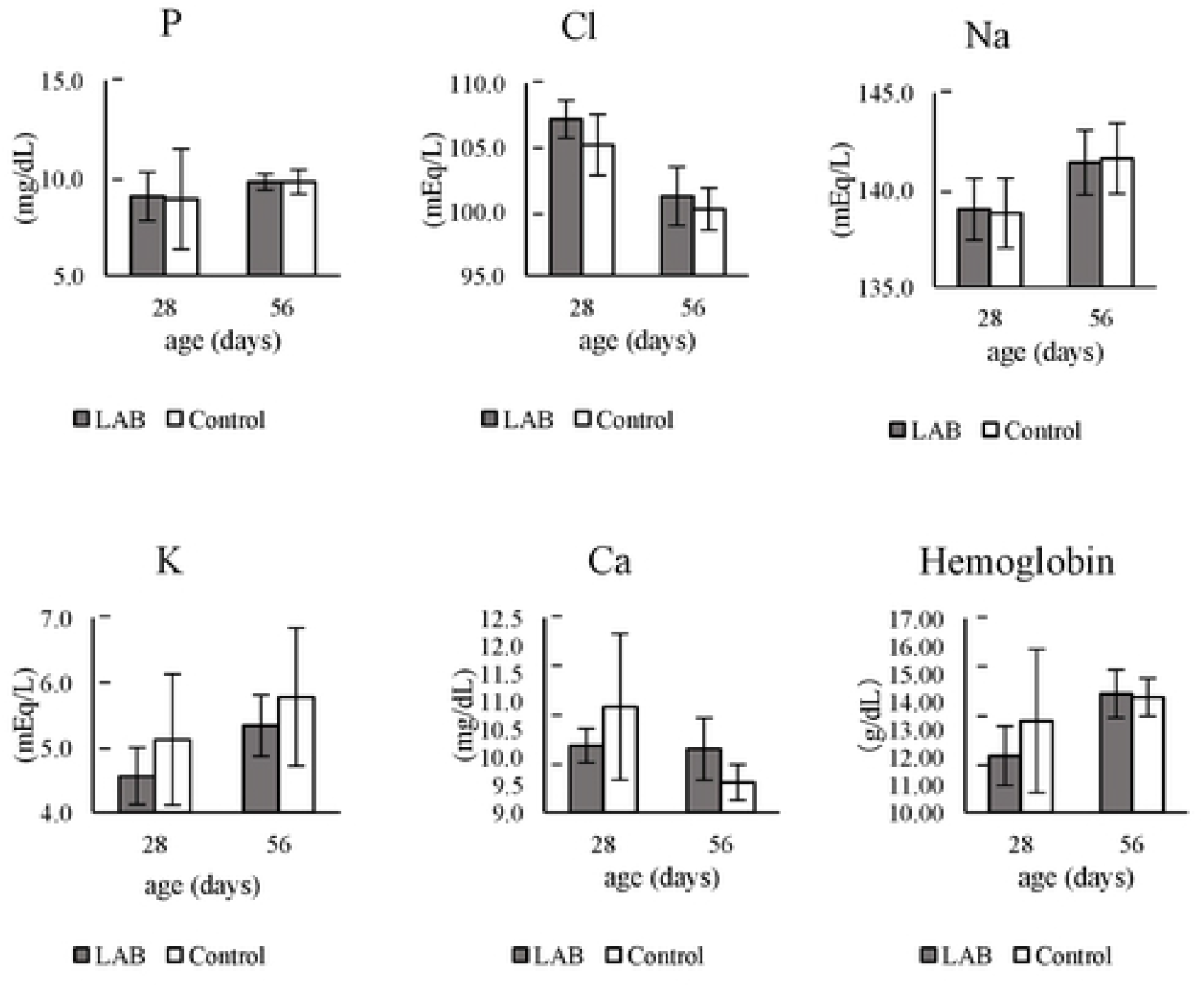
Blood biochemistry, blood cell counting of piglets in Test I. The gray bar indicates the average value of piglets A (LAB group (n=5) in Fig. 1), and the white bar indicates the average value of piglets D (Control group (n=5) in Fig. 1). The data is expressed as the mean ± standard deviation (SD).

**Table S1.** Composition of homemade diet. The composition of a homemade diet is shown as a percent. The homemade diet given to the piglets in Test I and Test II gradually increased from birth to 56 days of age.

| ingredients | % |
| --- | --- |
| corn | 33.0 |
| wheat | 20.1 |
| roasted soybean powder | 5.4 |
| soybean meal | 7.7 |
| wheat powder | 17.5 |
| skimmed milk powder | 10.3 |
| bran | 2.1 |
| vegetable oil | 1.0 |
| salt | 0.2 |
| calcium phosphate | 1.6 |
| calcium carbonate | 0.2 |
| minerals | 0.2 |
| Vitamin A,D,E complex | 0.1 |
| Vitamin B complex | 0.1 |
| L-lysine HCl | 0.4 |
| DL-methionine | 0.1 |
| total | 100.0 |

**Table S2.**
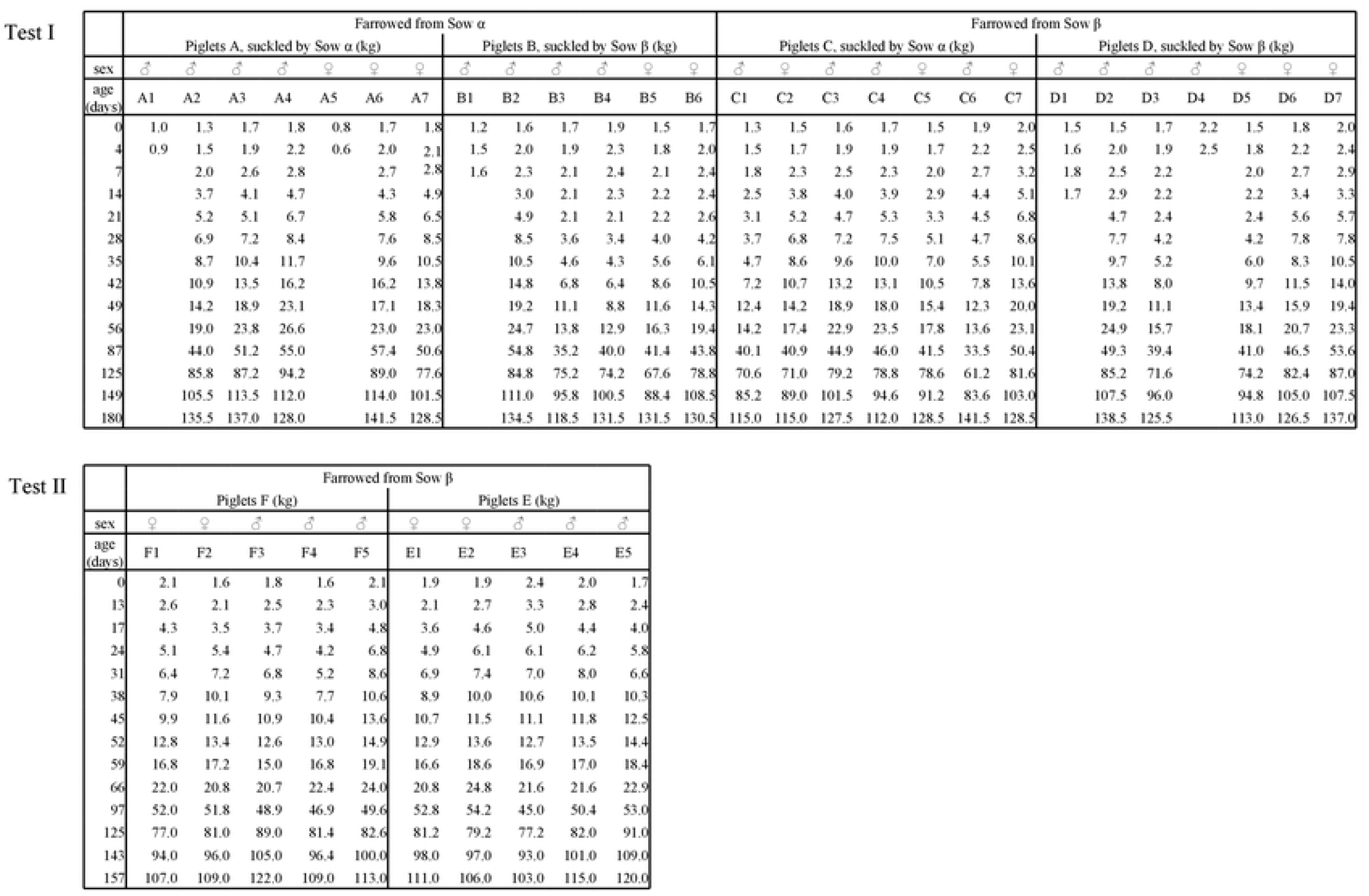
Body weight changes of piglets in Test I and Test II. Weighing results for individual piglets in Tests I and II are shown. The weight record for piglets that died during the rearing process remains blank.

## Notes

### Competing Interest Statement

The authors have declared no competing interest.

